# Rapid coordination of followership and leadership roles in homing pigeons navigating with unfamiliar partners

**DOI:** 10.64898/2026.07.06.736763

**Authors:** Joe Morford, Patrick J. Lewin, Lucy Larkman, Gayatri Kumar, Joy W. Kinuthia, Takao Sasaki, Richard P. Mann, Christopher Krupenye, Dora Biro

**Affiliations:** Department of Brain and Cognitive Sciences, University of Rochester, Rochester, New York, USA; Department of Biology, Oxford University, Oxford, UK; Department of Statistics, School of Mathematics, University of Leeds, Leeds, UK; Department of Psychological & Brain Sciences, Johns Hopkins University, Baltimore, Maryland, USA

**Keywords:** Collective movement, Coordination, Information transfer, Followership, Leadership

## Abstract

Collective movement requires coordination between individuals, yet how this emerges during early interactions remains poorly understood. We investigated how partner familiarity influences coordination, leader-follower dynamics, and learning in homing pigeon pairs navigating from novel sites. Birds were released repeatedly with either familiar or unfamiliar partners, followed by solo releases to assess learning. By quantifying bidirectional information flow, we found familiarity influenced information-transfer dynamics during the first release: familiar pairs exhibited more asymmetric information transfer, likely reflecting established leader-follower relationships, whereas unfamiliar pairs showed more symmetric exchange. These differences disappeared after one release. Conversely, familiarity had little effect on cohesion or navigational performance. There was some evidence for an influence on learning: birds from familiar pairings had higher homing efficiency on a subsequent solo release. Finally, across partnerships, followership was more predictable than leadership with respect to individual identity and flight speed, indicating stable variation in individuals’ tendency to follow rather than lead. This suggests that a shift in emphasis from leadership to followership might enhance our understanding of collective decision-making dynamics. Our results demonstrate how flight partners rapidly coordinate, producing limited downstream effects on navigation and learning, with implications for many animals that travel in fission-fusion transitory collectives.

## Introduction

Many of the tasks that animals face daily to survive, acquire resources, and reproduce, involve collective action with social partners within groups. These range from tasks involving joint, coordinated action, such as collective movement, to those requiring differentiated and coordinated complementary roles, for example biparental offspring provisioning or collaborative hunting. Considerable progress has been made in identifying the mechanisms that underpin such behaviours (Bailey et al., 2013; Baldan & van Loon, 2022; Couzin et al., 2005; Couzin et al., 2002; Gillies et al., 2022; Halliwell et al., 2022; Harcourt et al., 2009; Harcourt et al., 2010; Pettit et al., 2013; Rands et al., 2003; Wu et al., 2024); however, the role of learning in establishing, stabilising and refining coordination between individuals has received comparatively less empirical attention (Biro et al., 2016; Collet et al., 2023).

One of the most widespread collective coordination problems across animal taxa is the guidance of collective movement. This challenge arises across a wide range of scales and contexts, from localised predator avoidance and foraging to homing and long-distance migration (Berdahl et al., 2018). Homing pigeons (*Columba livia*) provide a powerful model system for studying the interaction between learning and collective decision-making during collective movement. When displaced from distant sites, pigeons typically return to their home loft in cohesive flocks, and their trajectories can be tracked simultaneously at high resolution using GPS devices. Repeated releases from the same site lead to marked improvements in navigational efficiency through learning and the development of stable, idiosyncratic routes, both in individuals (Biro et al., 2004; Guilford & Biro, 2014; Meade et al., 2005) and in flocks (Flack, Freeman, et al., 2013; Sasaki & Biro, 2017).

Furthermore, flocks show some differentiation of individual roles during homing, with decision-making hierarchies determining which birds lead and follow during homing (Flack, Akos, et al., 2013; Nagy et al., 2010; Nagy et al., 2013). These hierarchies have been inferred by quantifying how directional changes initiated by each bird propagate through the flock to other birds with sub-second delays, typically using correlation-based measures (Nagy et al., 2010). More recent work has extended this approach by measuring transfer entropy to quantify information transfer between birds, generating pairwise estimates of information flow that allow both the strength and asymmetry of leader–follower relationships to be quantified between individuals (Valentini et al., 2021). Leadership hierarchies are not related to social dominance relationships between birds, but instead are associated with, and likely mediated by, consistent spatial positioning, with leaders tending to occupy more frontal positions (Nagy et al., 2010; Valentini et al., 2021), and are predicted by individual traits such as flight speed and personality (Pettit et al., 2015; Pettit et al., 2013; Sasaki et al., 2018). However, it remains unclear how leadership and followership roles emerge and stabilise when individuals first navigate together, and whether learning is required to coordinate these differentiated roles. This question is particularly relevant when animals travel with unfamiliar partners, a situation that is common in animals that organise into fission–fusion societies or transitory aggreagations with unstable group membership, for instance in many avian taxa on migration (Harcourt et al., 2010). Previous work has shown that leadership hierarchies in pigeon flocks are relatively stable once established (Flack, Akos, et al., 2013), but little is known about how these relationships form during early interactions, or how prior experience with specific partners influences the dynamics of coordination and information use during navigation.

Leadership of homing pigeon flocks has been shown to predict individual learning, with birds that occupy leadership roles during flock flights subsequently exhibiting straighter routes when tested alone (Pettit et al., 2015). This pattern is consistent with group-level processes influencing individual learning, but could also be generated by underlying traits, such as homing speed, motivation, or personality, that simultaneously affect both leadership and learning (Pettit et al., 2015; Sasaki et al., 2018). Further, the superficial similarity between solo birds and pairs in their gradual improvements in homing efficiency and increases in route fidelity with repeated releases at a site might suggest that group context has limited influence on learning (Flack, Freeman, et al., 2013; Sasaki & Biro, 2017). Hence, it remains unclear whether individuals within flocks learn as if in isolation, or how learning might be shaped by leadership dynamics, group context, and collective decision-making processes (Kao et al., 2014; Morford et al., 2022).

In this study, we used pairs of homing pigeons to examine how learning and collective decision-making dynamics unfold when individuals navigate with familiar versus unfamiliar partners. Birds were released in pairs three times from novel sites either with a familiar partner, with whom they had extensive prior joint flight experience, or with an unfamiliar partner, with whom they had not previously homed that year (hence, “unfamiliarity” here signifies lack of recent joint homing experience, rather than never having encountered each other previously). To assess individual learning, pigeons were also released solo after the first and last paired releases. The experiment was conducted at two sites, such that each bird experienced both treatments. Using this design, we tested whether partner familiarity influenced pair cohesion, how directional information transfer and leadership asymmetries emerged across repeated interactions, and whether early coordination dynamics shaped navigational performance in pairs and subsequent individual learning.

## Methods

### Study System

This study was conducted using 38 birds, aged between 2 and 9 years, from a captive population of homing pigeons at the John Krebs Field Station, Oxford, U.K. In March and April of 2024, pigeons underwent a pretraining procedure to familiarise them with the experimental procedures. During this period, the pigeons were released from four different sites, each located approximately 2km from their home loft in distinct compass directions. Each bird was released four times in flocks and then four times individually from each site. Additionally, pigeons were acclimated to wearing harnesses and carrying plasticine weights (~15g), which accounted for less than 5% of their body mass and matched the weight of the GPS devices used in the experimental trials (described below). GPS devices recorded positional data at a 1 Hz resolution (Mobile Action iGot-U GT120).

### Partner Familiarity Treatment

Pigeons were randomly assigned into pairs, and these pairs were trained together 35-40 times in repeated releases across two training sites, with approximately half of the releases at each site (training site 1: 51.7527, −1.4237, 245° degrees and 8.1km from home; training site 2: 51.8303, −1.3915, 315° degrees and 7.3km from home). The birds trained together at these sites are referred to as ‘familiar partners’ in this study.

14 pairs of pigeons completed the full training. However, 1 bird went missing after the end of training, such that 13 complete pairs were available for testing. The 26 birds were re-paired in 13 new pairs, such that each was assigned a new, ‘unfamiliar partner’: a bird with whom it had not been paired in 2024. These unfamiliar pairings were assigned randomly, whilst ensuring that every bird had a distinct unfamiliar partner.

### Testing Phase

Test releases took place at two new release sites, sequentially (all releases completed at test site 1 before releases at test site 2), neither of which the pigeons had previously visited (test site 1: 51.7333, −1.2732; 151° and 6.3km from home; test site 2: 51.8017, −1.2260; 72° and 6.6km from home).

At each site, pigeons underwent a pre-testing release at the site in four groups (of 6 or 7 birds) to reduce the chance of the birds getting lost in subsequent experimental releases, with individuals in different groups than their experimental partners. Following the pre-testing release, pigeons were released in pairs three times at the site, and additionally released solo twice, after the first and last paired releases. Each bird was released with its familiar partner at one of these sites and its unfamiliar partner at the other.

One pigeon failed to return on its second paired release at the first site (in the familiar partner treatment), and was replaced at the second site with the pigeon whose training partner had gone missing after the end of training to retain a total sample of 26 birds (13 pairs). Two birds failed to return from the second site, one on the first paired release, and one on the second paired release (both in the unfamiliar partner treatment).

### Data processing

GPS tracks were trimmed from the point at which they left 50m of the release site to the point at which they entered the 500m radius of the home loft. This removed the initial post-release escape response of the pigeons and the final section of homing in which the pigeons are within visual contact of their home loft. Portions of the track in which the bird was moving less than 30 kilometres per hour were removed to isolate tracks in which the bird was flying rather than sitting.

To identify cases in which the pigeons that had been released separately joined and flew together post-release, tracks from the same release were cross-checked. 9 tracks were then excluded on the basis of the birds joining together post-release.

### Information transfer and leadership

To examine information transfer between birds, we used transfer entropy to quantify the extent to which a bird’s directional changes could be predicted by those of its partner at the previous time-step, while accounting for autocorrelation in the focal bird’s trajectory, as in (Valentini et al., 2021). This approach allows information transfer to be quantified separately in each direction (information transfer from each bird, and to each bird). We used a binary encoding of point-to-point directional changes (clockwise vs anticlockwise) and a history length of 3 samples to account for autocorrelation in the receiver’s trajectory. This history length was chosen to best balance the benefit of capturing longer temporal dependencies against ensuring sufficient sampling of all possible history sequences (the number of which grows as a power-law of history length). We therefore retained tracks for analysis in which each possible history sequence contained, on average, at least 25 observations (at least 200 total observations before, or without, the birds splitting). This criterion was met by over 90% of eligible tracks (compared to only 49% if a history length of four had been used).

We quantified information transfer asymmetry by calculating the difference between the pairwise transfer entropy measures for a pair of birds.

### Analysis

In general, across analyses, we fit linear mixed-effects models and tested for reductions in model fit with likelihood ratio tests (LRTs) when dropping the interaction between treatment and release number, and subsequently dropping treatment from the model altogether. Pairwise differences between treatment groups within each paired release were assessed using estimated marginal means, with p-values based on asymptotic Wald z-tests.

#### Pair cohesion

To test whether partner familiarity influenced pair cohesion, we examined whether the experimental treatment predicted the probability of pair splitting. Splitting was defined as birds moving more than 150m apart and failing to re-establish proximity within this distance for the remainder of the flight. Splitting was modelled as a binary response using a binomial error distribution in a generalised multimember mixed-effects model: *Split (True/False) ~ Treatment * Paired_Release_Number + Site + (1 | Individual_ID) + (1 | Pair_ID)*. Paired_Release_Number was treated as a categorical variable. We analysed the mean inter-individual distance (log-transformed to meet the assumptions of linear modelling) between paired birds (excluding flight segments following a split) over the entirety of homing trajectories, and also in the initial decision-making section of homing trajectories (before leaving 2km of the release site for the final time, as in (Morford et al., 2024)). Finally, we analysed the log-transformed rate of swaps in the identity of the bird that was flying in front (excluding sections after a split). For each of these response variables, a single datapoint, averaged across the pair for each release, was used to avoid pseudoreplication in multimember generalised linear mixed-effects models: *Response ~ Treatment * Paired_Release_Number + Split + Site + (1 | Individual_ID) + (1 | Pair_ID)*.

We examined the relationship between pair cohesiveness and navigational performance by testing whether birds that split had a lower homing efficiency that those that remained cohesive. Homing efficiency was quantified using the Homing Efficiency Index (HEI; (distance to home of start of trajectory – distance to home of end of trajectory) / total distance moved), calculated from the point birds left the 2km vicinity of the release site for the final time. HEI was modelled with a linear mixed-effects model assuming a beta error distribution with a logit link function: *HEI ~ Split * Paired_Release_Number + Site + (1 | Individual_ID) + (1 | Pair_ID)*.

#### Information transfer

In order to test the influence of partner familiarity on the magnitude of information transfer between birds, we predicted transfer entropies (log-transformed to meet the assumptions of linear modelling given the right-skewed, strictly positive distribution) with a linear mixed-effects model: *log(Transfer Entropy i→j) ~ Treatment * Paired_Release_Number + Split + Site + (1 | Individual-i_ID) + (1 | Individual-j_ID)*. We further tested for differences in the two-dimensional distributions of pairwise transfer entropy values (to each bird and from each bird) on each release using Hotelling’s T^2^ tests, subsetting to include the bird in each pair for which the information transfer to the bird was greater than the information transfer from the bird to avoid pseudoreplication, and obtaining non-parametric p-values through permutation (n = 10,000). We examined asymmetry in information transfer with a multimember linear mixed-effects model*: log(Absolute Information Transfer Difference) ~ Treatment * Paired_Release_Number + Split + Site + (1 | Individual_ID) + (1 | Pair_ID)*.

#### Drivers of information transfer dynamics

To estimate the repeatability of the information transfer from each bird and to each bird, we subsetted the dataset to include birds that had at least 4 observations (as either the bird transmitting or receiving information) and fit a linear mixed-effects model including the identities of individuals transmitting and receiving information as random effects, bootstrapping with 1,000 iterations to generate confidence intervals for the repeatability measures. Further, we examined the difference between the repeatabilites associated with the individual identities (from individual versus to individual) using a simulation-based test of variance components: we repeatedly simulated data from the model (1,000 iterations) and refit it to each simulated dataset to obtain confidence intervals of the difference. Additionally, we tested whether solo flight speeds (in their first solo release) predicted information transfer in subsequent paired releases (averaged across the subsequent releases) using a mixed-effects model: *log(Transfer Entropy i→j) ~ Bird i Solo Flight Speed + Bird j Solo Flight Speed + Site + (1 | Individual-i_ID) + (1 | Individual-j_ID)*.

#### Navigational performance

We assessed treatment effects on pair navigational performance using three metrics: the HEI (calculated from the final departure from the release site), the Initial HEI (the efficiency of homeward approach prior to leaving the 2 km vicinity of the release site for the final time), and the Virtual Vanishing Time (VVT; time taken to leave the 2 km vicinity for the final time; log-transformed to improve normality). Analyses were restricted to pairs that did not split during the relevant track sections. A single point averaged across pairs of birds for each release was used in a multimember linear mixed-effects model *Response ~ Treatment × Paired_Release_Number + Site + (1 | Individual_ID) + (1 | Pair_ID)*, using Gaussian error distributions in each case to incorporate multimember modelling and negative values of efficiency indices.

#### Navigational learning

Finally, we examined route fidelity and homing efficiency of solo trajectories relative to the previous paired release. Route fidelity was quantified as the mean distance between a pigeon’s solo homing trajectory (after leaving the release-site vicinity) and its previous paired trajectory; cases where pairs had split were excluded, and values were log-transformed to meet model assumptions. Route fidelity was analysed using a linear mixed-effects model (*Route fidelity ~ Treatment × Solo_Release_Number + Site + (1 | Individual_ID) + (1 | Pair_ID)*). Solo homing efficiency was calculated using the HEI from the point of final departure from the release-site vicinity and modelled with a beta-distributed mixed-effects model with a logit link (*HEI ~ Treatment × Solo_Release_Number + Split_Previous_Release + Site + (1 | Individual_ID) + (1 | Pair_ID)*). To assess the role of information transfer dynamics, we tested whether pairwise transfer entropies from the previous paired release predicted route fidelity and homing efficiency using linear mixed-effects models including log-transformed transfer entropies (both from bird and to bird), Solo_Release_Number, and Site, with Individual-A_ID as a random effect.

## Results

### Pair cohesion

We observed a reasonably high rate of birds splitting from their flight partners, especially on their first paired release (observed in 11 out of 25 unique pairs), but birds were equally likely to split from highly familiar or unfamiliar flight partners (Fig. 1A; linear multimember mixed-effects model: 71 observations/25 pairs/27 birds; likelihood ratio tests: dropping interaction with release number, p = 0.735; dropping treatment altogether, p = 0.748). Splitting was associated with reduced homing efficiency across releases (Fig. 1B; generalised linear mixed-effects model with 142 observations/25 pairs/27 birds; LRTs: dropping interaction with release number, p = 0.538; dropping splitting altogether, p < 0.0001). However, it is difficult to definitively determine whether this relationship is causal, and whether birds that split would have achieved higher efficiency had they remained together.

**Figure 1.**
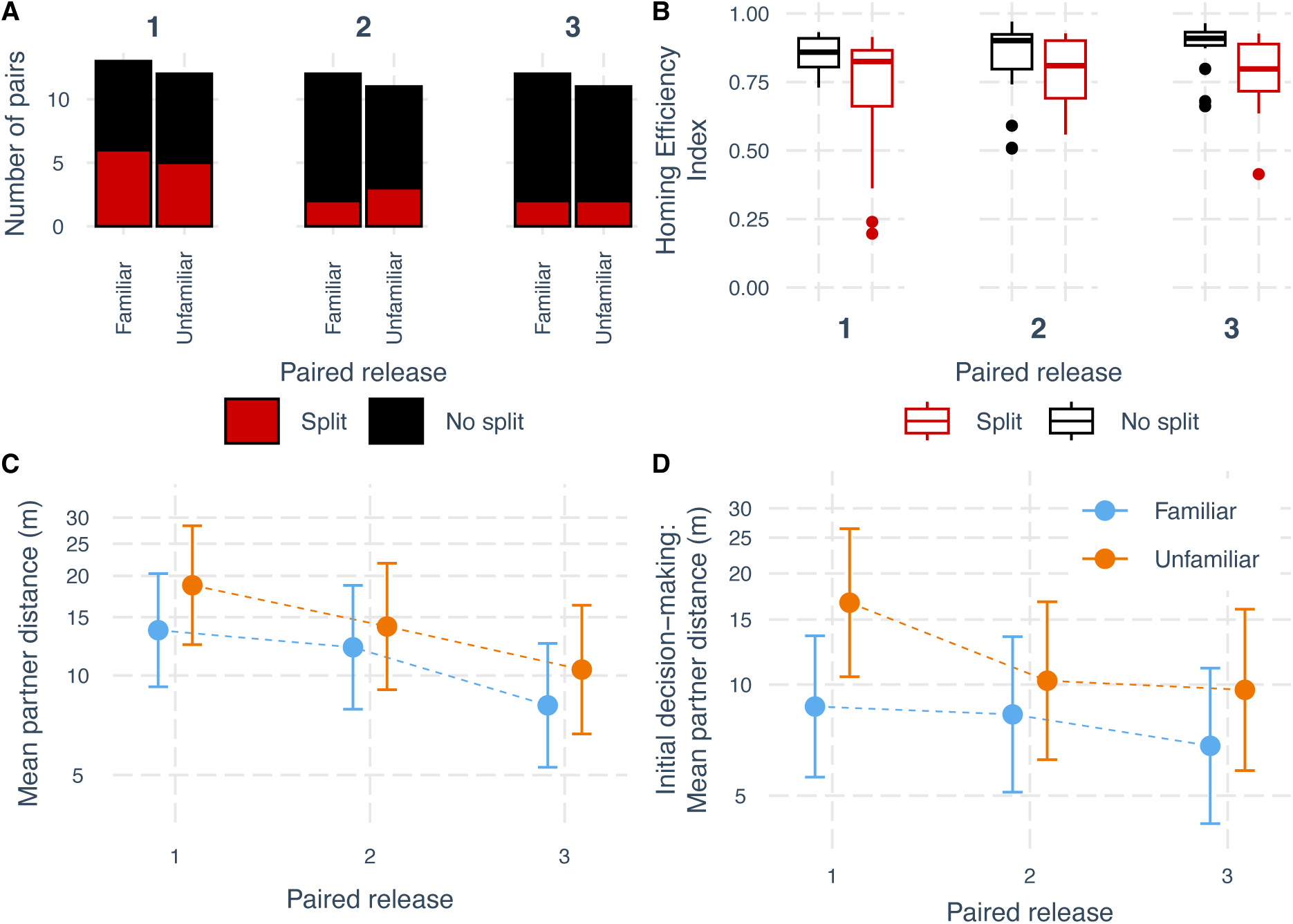
Transient effect of partner familiarity on pair cohesion. (A) Frequencies of pairs splitting across releases for familiar and unfamiliar pairings. (B) Homing efficiency (HEI) of birds that remained cohesive versus those that split. (C) Mean inter-individual distance between paired birds across entire homing trajectories. (D) Mean inter-individual distance during the initial decision-making phase (before leaving the 2 km vicinity of the release site). Points represent estimated marginal means ± 95% confidence intervals from full models.

To further examine the influence of partner familiarity on pair cohesion, we tested for differences in mean inter-individual distance during homing. We found no significant difference in the mean distance of familiar versus unfamiliar partners across paired releases (Fig. 1C; 71 observations/25 pairs/27 birds; LRTs: dropping interaction with release number, p = 0.805; dropping treatment altogether, p = 0.136). In contrast, examining inter-individual distances in the initial decision-making section of homing tracks (before leaving the 2km vicinity of the release site for the final time) revealed differences in pair cohesion between familiar and unfamiliar pairings across releases (Fig. 1D; multimember mixed-effects model with 71 observations/25 pairs/27 birds; LRTs: dropping interaction, p = 0.441; dropping treatment, p = 0.011). This was most pronounced on the first release, with familiar partners flying significantly closer than unfamiliar partners (p = 0.021), whereas there was no significant difference on the second or third release (p = 0.453 and p = 0.216). This pattern may reflect the process by which birds learned how to coordinate their movements with an unfamiliar flight partner in the first part of their first release together. Finally, we found no differences in the rate of swaps in the identity of the bird flying in front between treatments across paired releases (71 observations/25 pairs/27 birds; LRTs: dropping interaction, p = 0.649; dropping treatment: p = 0.541). Together, these results suggest that early differences between familiar and unfamiliar pairings in cohesion are relatively transient and hence partner familiarity has limited influence on overall pair cohesion.

### Information transfer

We used a transfer entropy measure to quantify the bidirectional information transfer between pairs of birds during their homing journeys. We found no differences in the magnitude of information transfer between treatment groups across paired releases (Fig. 2A; 128 observations/27 birds; LRTs: dropping interaction, p = 0.527; dropping treatment, p = 0.430). Additionally, we found no difference in the magnitude of information transfer between those pairs that subsequently split versus those that remained cohesive (LRT: p = 0.295).

**Figure 2.**
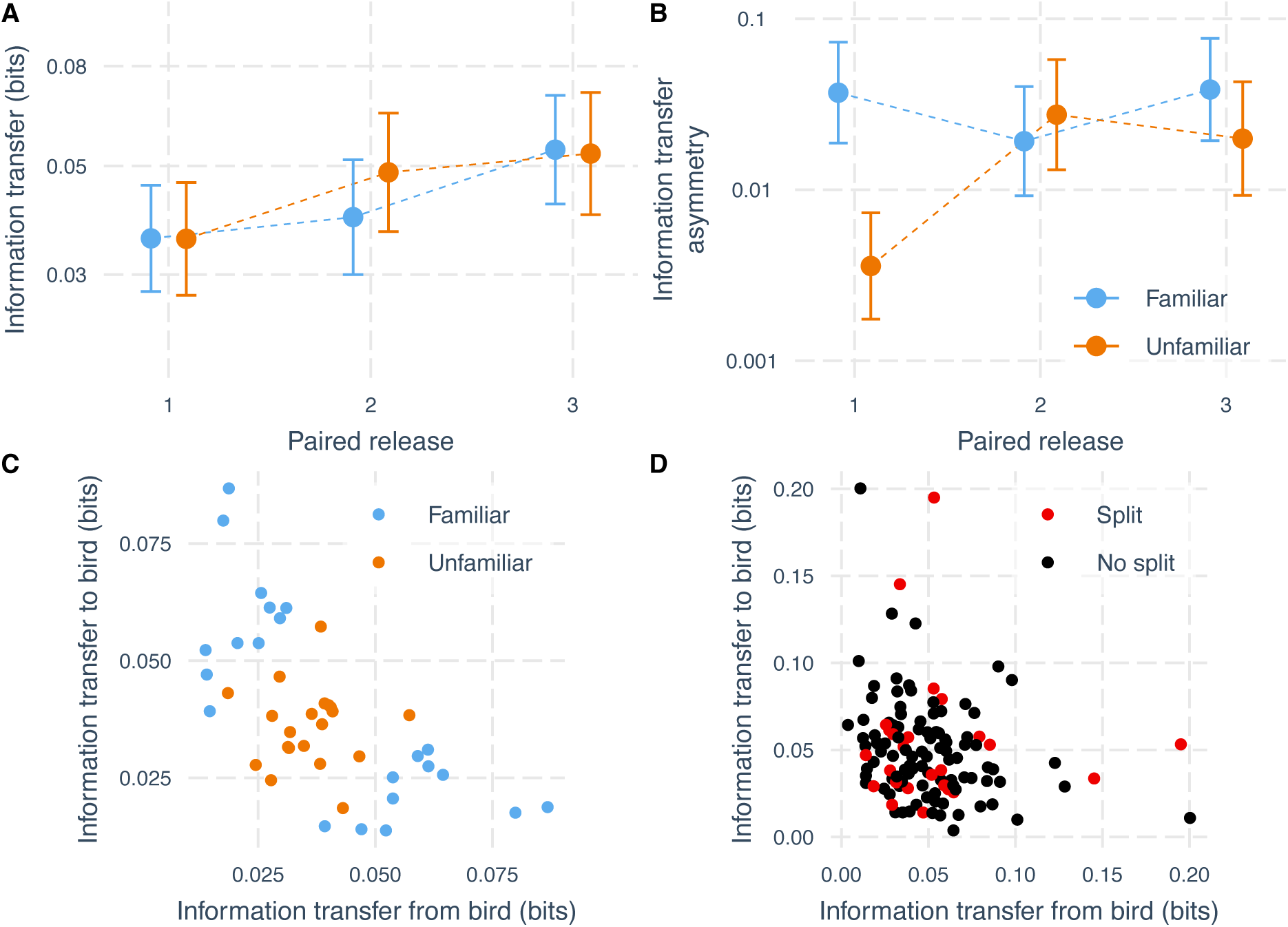
Partner familiarity shapes early information-transfer dynamics. (A) Magnitude of pairwise transfer entropy between birds across paired releases. Points represent estimated marginal means ± 95% confidence intervals from full models. (B) Asymmetry in information transfer (absolute difference between directions) for familiar and unfamiliar pairs. (C) Two-dimensional distributions of directional transfer entropy values for familiar and unfamiliar pairs on the first paired release. (D) Corresponding distributions for pairs that later split versus those that remained cohesive, across all three paired releases.

However, examining the asymmetry in paired information transfer, quantified as the difference in the magnitude of information transfer in each direction between a pair of birds, revealed differences between familiar and unfamiliar pairings, with a significant interactive effect of treatment and release number (Fig. 2B; 64 observations/25 pairs/27 birds; LRT: dropping interaction, p = 0.0002). We found higher asymmetry between familiar pairings than between unfamiliar pairings (p < 0.0001) in the first paired release, but no difference in the second (p = 0.468) and or third (p = 0.176) releases. Further, we found that asymmetry was significantly smaller for unfamiliar pairings in the first paired release compared to the second (p = 0.0001) or third (p = 0.001) paired releases (no difference between second and third paired releases: p = 0.510). Conversely, there was no difference in the asymmetry strength of familiar birds between any of the paired releases (first-second: p = 0.172; second-third: p = 0.130; first-third: p = 0.927). To support this evidence, we performed Hotelling’s T^2^ tests to examine whether distributions of pairwise transfer entropies differed between treatment groups on each release. We found that familiar and unfamiliar birds had significantly different distributions of pairwise transfer entropies in the first paired release (Fig. 2C; T^2^ = 19.7, p < 0.0001, df = (2, 18)), but no evidence that they were different on the second (T^2^ = 1.14, p = 0.350, df = (2, 18)) or third paired releases (T^2^ = 0.537, p = 0.600, df = (2, 19)). These findings indicate that partner familiarity primarily influences the early dynamics of information transfer, with unfamiliar pairs initially exhibiting more symmetrical interactions before rapidly converging on patterns similar to familiar pairs.

We found no evidence that pairs that ultimately split versus those that remained cohesive had distinct pairwise transfer entropy distributions in the first two paired releases (release 1: T^2^ = 0.280, p = 0.754, df = (2, 18); release 2: T^2^ = 0.005, p = 0.994, df = (2, 18)). However, we did find a marginally significant difference in these distributions in the third paired release (T^2^ = 4.40, p = 0.038, df = (2, 19)). This perhaps indicates a difference in the nature of the information transfer between pairs with some experience that remain cohesive compared to those that split. The distribution of pairwise transfer entropies for pairs that split and those that remained cohesive, across releases, is shown in Figure 2D. Further work could explore whether splits occurring in later releases are preceded by identifiable changes in information transfer dynamics.

### Drivers of information transfer dynamics

The identity of the individual transmitting information did not explain any variation in the magnitude of information transfer (Fig. 3A) and hence had a corresponding repeatability of 0%. Conversely, we found that the magnitude of information transfer had a repeatability of 30.1% for the individual receiving information (Fig. 3B); this was significantly different from 0 (Likelihood Ratio Test: p < 0.01). Hence, variation in information transfer was more strongly associated with the identity of the receiving bird than the transmitting bird; however, the difference in repeatabilites was not significant with simulation demonstrating that the confidence interval of the difference overlapped with 0 (−0.134, 0.483). We examined the relationship between mean front–back distance within a pair and directional information transfer using local non-linear regression (loess; Fig. 3C). Consistent with previous work (Valentini et al., 2021), information transfer was substantially greater from the bird in front to the bird behind.

**Figure 3.**
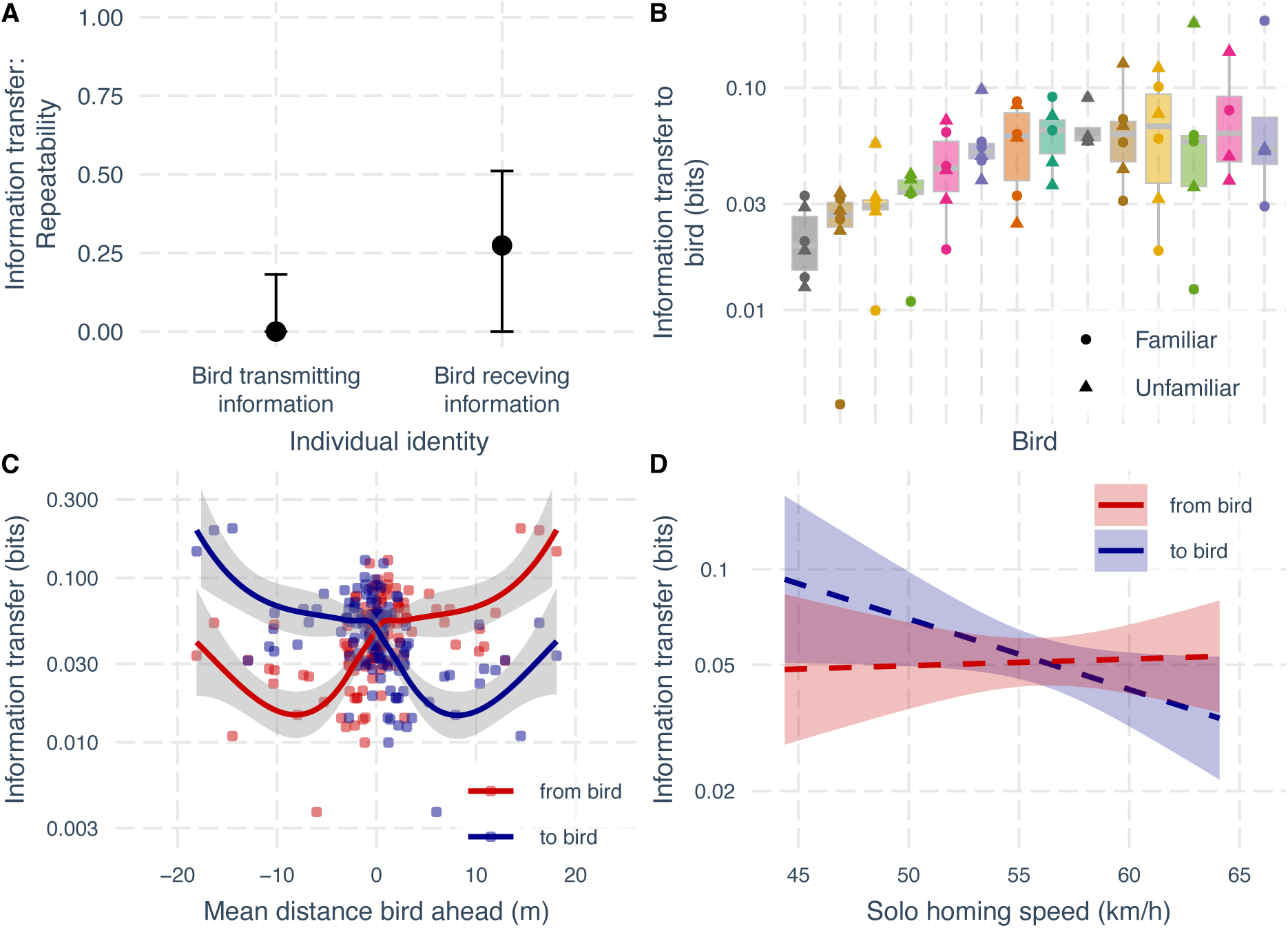
Drivers of information transfer dynamics. (A) Repeatability of information transfer magnitude between birds as function of the identity of the bird transmitting information and receiving information. Information transfer was significantly repeatable for the birds receiving information, but not for the birds transmitting information. (B) Information transfer received by each bird across releases and partners (from subset dataset including birds with at least four observations). (C) Relationship between mean front–back distance and directional information transfer during paired flights. Curves show local non-linear regression fits (loess). Information transfer is substantially greater from the bird in front to the bird behind than in the reverse direction, consistent with previous findings. (D) The significant negative relationship between information transfer from bird A to bird B during paired flights and the solo flight speed of the bird receiving information: bird B. The line shows the model-predicted relationship with 95% confidence intervals.

Based on previous work showing that solo flight speeds predict leadership hierarchies in subsequent flock releases (Pettit et al., 2015), we expected a positive relationship between the solo flight speed of a bird and the magnitude of information transfer from the bird, but a negative relationship with the magnitude of information transfer to the bird. We did not find the predicted positive relationship with the magnitude of information transfer from the bird (estimate = 0.005, p = 0.816), but did find the expected negative relationship with the magnitude of information transfer to the bird (Fig. 3D; estimate = −0.052, p = 0.028). Therefore, our results support previous work by showing that solo flight speeds can predict decision-making dynamics, but again suggest that followership may be more repeatable and predictable than leadership.

### Navigational performance

Examining overall navigational performance, we found no effect of treatment on the Homing Efficiency Index of paired flights (Fig. 4A, 51 observations/23 pairs/27 birds; LRTs: dropping interaction with release number, p = 0.799; dropping treatment, p = 0.259). Next, we examined two metrics of initial navigational performance, the Initial Homing Efficiency Index and the Virtual Vanishing Time (the time taken to leave the vicinity of the release site). We found some evidence for an interaction between treatment and release number on the Initial Homing Efficiency Index (63 observations/25 pairs/27 birds; LRT: p = 0.037). This may indicate that initial homing efficiency increases more quickly with experience in unfamiliar pairings than in familiar pairings, as can be observed in Figure 4B. However, we found no significant difference between treatments on any of the three paired releases individually (release 1: p = 0.277; 2: p = 0.503; 3: p = 0.135). Further, we found no interactive or additive effect of treatment on the Virtual Vanishing Time (Fig. 4C; 63 observations/25 pairs/27 birds; LRTs: p = 0.127, p = 0.983). Taken together, these results provide little evidence that differences in information transfer or partner familiarity translate into meaningful differences in navigational performance during paired flights. While we did find evidence for a faster improvement in one of the metrics of initial navigational performance for unfamiliar pairings, this effect was unexpected and further work is required to establish the robustness and biological relevance of this effect.

**Figure 4.**
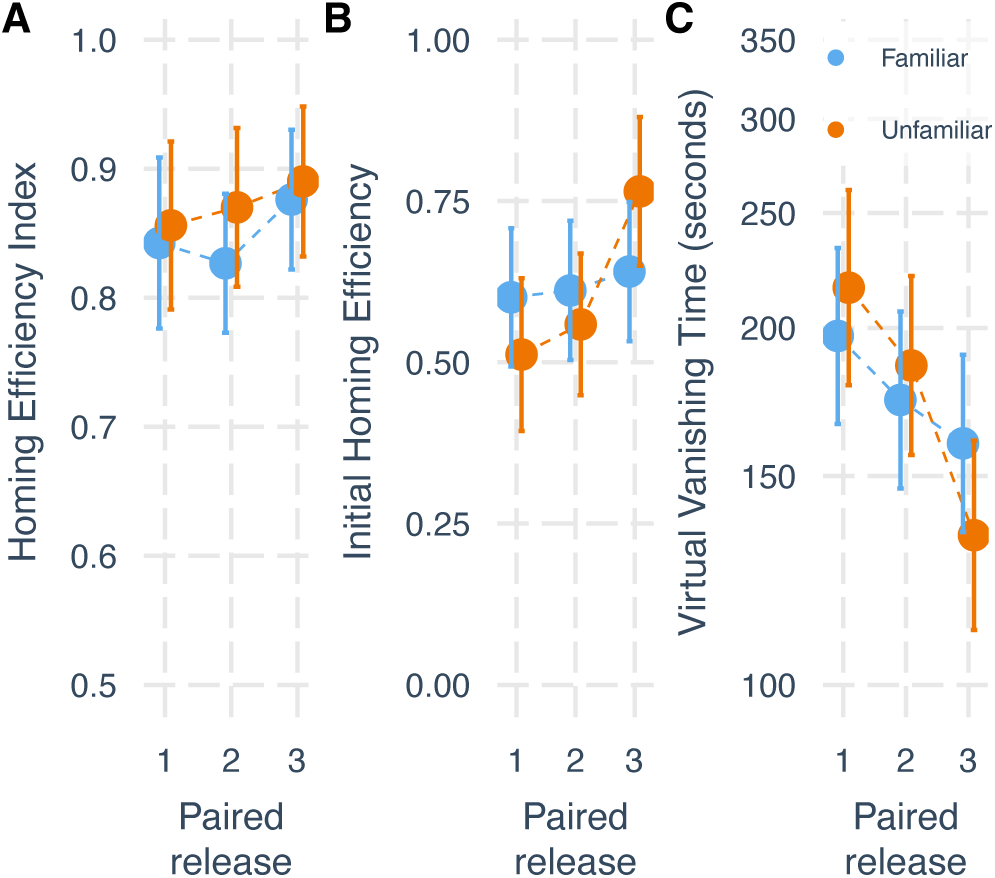
Limited evidence that partner familiarity affects navigational performance. (A) Homing Efficiency Index across paired releases. (B) Initial Homing Efficiency Index prior to leaving the 2 km vicinity of the release site. (C) Virtual Vanishing Time (time to leave the 2 km vicinity). Points represent estimated marginal means ± 95% confidence intervals from full models. There was a marginally significant interative effect between treatment and release on Initial Homing Efficiency, but no significant effect of treatment on any single release. There was no evidence of an effect of treatment on the other metrics of navigational performance.

### Navigational learning

We examined the influence of partner familiarity on the individual learning of navigational routes by examining how close birds flew to their paired routes and how efficiently they returned home when subsequently tested solo. The individual routes home taken by birds in the first solo releases at each site are shown in Figures 5A and 5B. We found no evidence of an effect of treatment on the mean distance of solo routes to the previous paired routes (Fig. 5C; LRTs: dropping interaction with release number, p = 0.711; dropping treatment, p = 0.198). Conversely, examining the homing efficiency of solo routes, we found some evidence for an effect of treatment, with a marginally non-significant drop in model fit when dropping treatment from the model (LRTs: dropping interaction: p = 0.532; dropping treatment, p = 0.059). Further, we found a significantly higher homing efficiency on the first paired release (p = 0.045) in birds that had been paired with a familiar partner, but no difference on their second paired release (p = 0.317). This suggests there is a weak effect of partner familiarity on navigational learning, with birds in familiar pairings initially learning more efficient routes than birds in unfamiliar pairings; this is consistent with the idea that attention to social cues may initially compete with attention to navigational cues in unfamiliar pairings. Finally, we tested whether pairwise information transfer values predicted both of these metrics of individual learning. In each case, fitting linear mixed-effects models to the mean distance of solo routes to their previous paired route (59 observations/22 pairs/26 birds) and to the homing efficiency of solo routes (78 observations/24 pairs/26 birds), we found that neither the magnitude of information transfer from a bird (p = 0.813 and p = 0.301, in the two models, respectively) nor to a bird (p = 0.289; p = 0.357) significantly predicted individual learning. Overall, these findings suggest that any effect of partner familiarity on individual learning is weak and short-lived, and that variation in information transfer during paired flights does not strongly predict subsequent solo performance.

**Figure 5.**
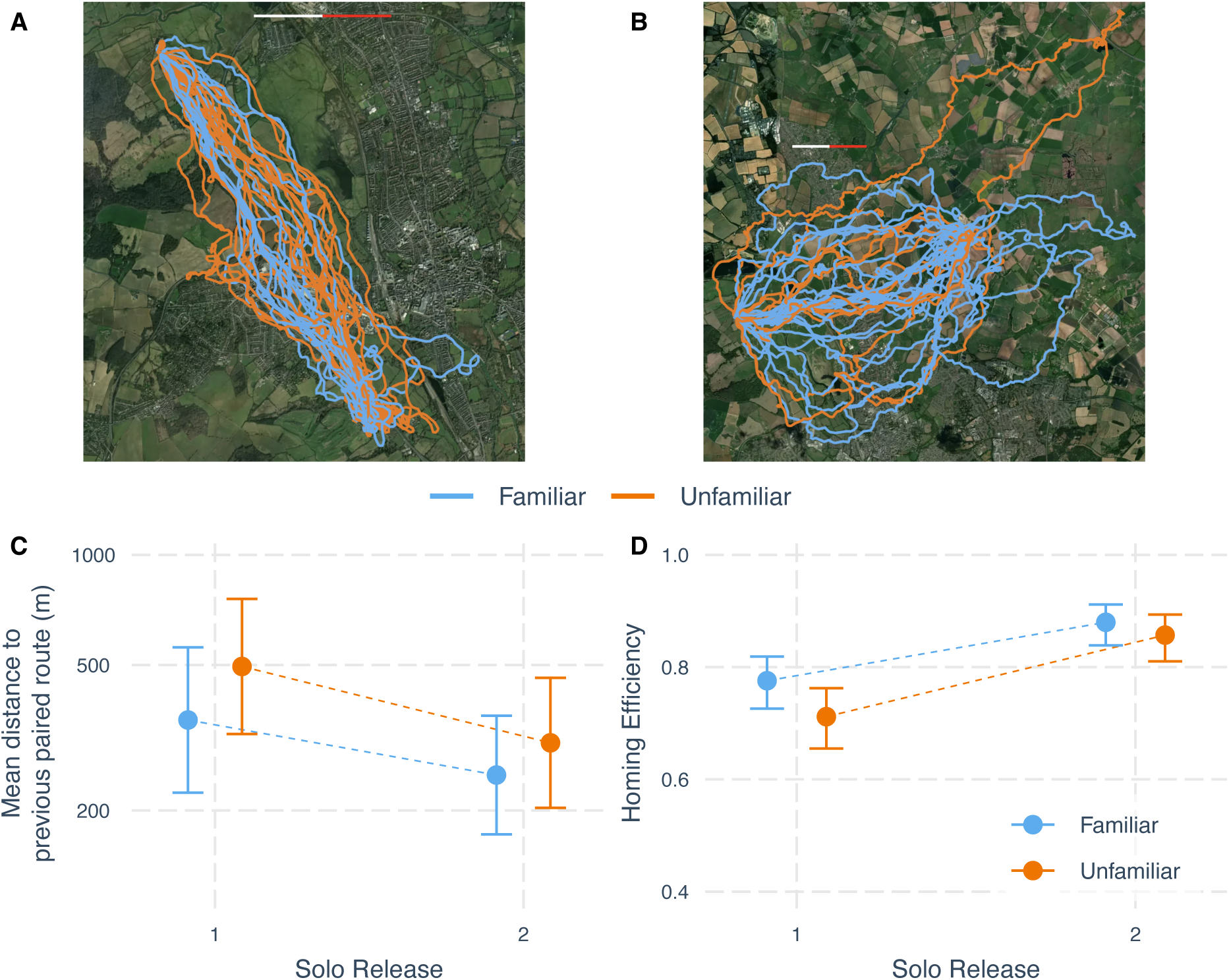
Transient effect of partner familiarity on individual learning. (A–B) Solo homing trajectories on the first solo release at each release site following familiar and unfamiliar paired releases. A kiolmetre scale bar is included (of total length =2km). The map was generated by the authors using Plotly (Python, version 3.11.7; https://plotly.com/python/) with Mapbox satellite basemap tiles (https://www.mapbox.com/). Map data, Mapbox, OpenStreetMap contributors. (C) Route fidelity, measured as the mean distance between solo trajectories and previous paired routes. (D) Homing efficiency of solo releases. Points show estimated marginal means ± 95% confidence intervals from full models. There was no evidence of an effect of treatment on route fidelity, but some evidence on an effect on solo Homing Efficiency, with a significant difference in Homing Efficiency between treatments on the first paired release, but no difference on the second release.

## Discussion

We used pairs of homing pigeons to examine how learning and collective decision-making dynamics unfold when individuals navigate with familiar versus unfamiliar partners from novel release sites. We found that partner familiarity influenced how coordination emerged between individuals, affecting early information transfer dynamics but having limited effects on cohesion and navigational performance.

The most pronounced differences between familiar and unfamiliar pairings emerged in the dynamics of information transfer. On the first paired release, unfamiliar pairs exhibited intermediate and relatively symmetrical levels of information transfer, whereas familiar pairs showed more asymmetric information flow between partners. Quantifying this asymmetry revealed substantially greater differences between flight partners in familiar pairs than in unfamiliar pairs on the first release. However, unfamiliar pairs reached levels of asymmetry similar to those of familiar pairs from the second paired release onwards. This temporal pattern indicates that unfamiliar pairs may initially engage in a process of coordinating or negotiating leadership roles, before rapidly adopting more stable roles thereafter. Further, these findings suggest that leadership dynamics within pairs may not be fixed, but instead rapidly negotiated during initial interactions. The absence of such a coordination period in familiar pairs supports previous work (Pettit et al., 2015) showing that leadership relationships are not dependent on site-specific knowledge, but instead generalise across contexts, including homing from novel release sites.

Using pairwise information transfer measures further allowed us to distinguish leadership, the extent to which a partner responds to an individual, from followership, the extent to which an individual responds to its partner (Nakayama et al., 2013). Notably, variation in information transfer was more strongly associated with the identity of the bird receiving information than the bird transmitting information, indicating that followership was more repeatable than leadership (albeit non-significantly). This interpretation is supported by the significant negative relationship between the magnitude of information received by an individual and its speed on a previous solo release, whereas no relationship was detected for information transmitted. Together, these findings suggest that individual differences in responsiveness to social information may be more stable than differences in influence over others, reframing previous work that linked individual traits such as flight speed or personality to leadership-followership dynamics without explicitly separating these roles (Pettit et al., 2015; Pettit et al., 2013; Sasaki et al., 2018). Further, an emphasis on followership rather than leadership may help to clarify what predicts the success or failure of the initiation of (changes in) collective movement (King, 2010; Petit & Bon, 2010).

The use of transfer entropy to quantify information transfer between birds builds on recent work in homing pigeons (Valentini et al., 2021) and offers several advantages in certain conditions over correlation-based measures used elsewhere to infer leadership (Nagy et al., 2010). In particular, transfer entropy provides directional, pairwise estimates of information flow, enabling the strength and asymmetry of leadership–followership relationships to be quantified between individuals. We demonstrate that this approach can be robustly applied to 1Hz data by replicating expected relationships between front–back distance and information transfer (Fig. 3C, like in (Valentini et al., 2021)), and by showing significant repeatability across independent measures from the same individuals. These results support the utility of transfer entropy for investigating coordination dynamics in small groups and suggest it could be extended to larger flocks. More generally, this highlights the value of information-theoretic approaches for uncovering the mechanisms underlying collective behaviour by quantifying the strength of leadership and followership between group members, and the distribution of influence on collective decision-making within groups.

Despite the pronounced differences in information transfer dynamics observed on the first paired release, we found limited evidence that these translated into differences in pair cohesion or navigational performance. Weak cohesion between pairs of pigeons, reflected in relatively high rates of splitting, are often observed on first releases at novel sites, before splitting rates decrease rapidly as birds are successively released (Flack, Freeman, et al., 2013). Whether this early decline reflects changes in social dynamics or reductions in navigational conflict as birds improve their homing performance remains unclear. Here, we have shown that birds were equally likely to split from highly familiar flight partners, with which they have been trained 35-40 times, as from unfamiliar flight partners. Together with the finding that any effects of familiarity on pair cohesion were limited to an initial coordination phase during the first paired release, this indicates that variation in pair cohesion may be more parsimoniously explained by navigational conflict between birds (as in (Biro et al., 2006)), rather than by social instability or unfamiliarity. Further, this suggests that better aligned directional preferences rather than greater familiarity might explain why, in previous work, pairs of pigeons trained together have been found to fly closer together while homing in larger flocks from the same release site (Flack, Freeman, et al., 2013). More broadly, this supports the idea that cohesion may often depend on the levels of conflict in information or goals between group members, rather than the strength of social relationships between them. Compared to the results on the development of information transfer dynamics and coordination above, these findings on group cohesion are more consistent with self-organisation models that typically do not incorporate explicit social relationships, instead generating collective patterns from simple local interaction rules (Ballerini et al., 2008; Couzin et al., 2005; Couzin et al., 2002; Herbert-Read et al., 2011; Katz et al., 2011).

Finally, we found some evidence that partner familiarity influenced individual navigational learning. Birds that had flown with a familiar partner returned more efficiently on their first solo test than birds previously paired with an unfamiliar partner. This pattern is consistent with the idea that attention to social cues may initially compete with attention to navigational cues in unfamiliar pairings, and aligns with previous suggestions that leaders may learn more quickly than followers because they pay more attention to spatial rather than social cues (Burt de Perera & Guilford, 1999; Kano et al., 2021; Lamprecht, 1991; Pettit et al., 2015). The more pronounced effect in the first solo test mirrors the timescale over which information transfer dynamics differed between familiar and unfamiliar pairs, suggesting that this transient coordination phase may generate short-lived competition between attention paid to social and navigational cues. More broadly, this highlights a general trade-off in collective systems, whereby the processes that enable individuals to coordinate with others may temporarily constrain their ability to acquire independent information. In contrast, we found no evidence that variation in pairwise information transfer predicted subsequent solo homing efficiency, nor that partner familiarity or information transfer influenced route fidelity. This somewhat contrasts a previous study showing that leaders learn more efficient routes after learning in flocks of 10 birds (Pettit et al., 2015), possibly because any such effects are much smaller or absent in pairs compared to larger flocks. Taken together, these findings suggest that any effect of partner familiarity on navigational learning is relatively weak and transient, and that variation in leadership dynamics in pairs does not strongly shape individual learning outcomes, perhaps contrasting stronger influence of coordination and leadership dynamics on individual learning in larger flocks.

These findings demonstrate how birds are able to rapidly coordinate with unfamiliar flight partners, with implications for the coordination dynamics and navigation during collective movement, especially for animals that often travel with unfamiliar partners in fission-fusion societies or in transitory aggregations (Harcourt et al., 2010). Similar coordination processes may occur in larger groups when spatial structure produces predictable interaction patterns that similarly facilitate the learning and stabilisation of coordination between adjacent group members (Flack, Freeman, et al., 2013; Nagy et al., 2010). However, moving in larger groups likely provides greater coordination challenges than flying in pairs, potentially extending the the process of coordination and negotiation of leadership roles and producing more pronounced downstream effects on navigational performance and learning. Further work could clarify these possibilities by extending our approach to examine coordination in larger flocks of homing pigeons and in other animal groups.

## Conclusions

In conclusion, our results demonstrate that homing pigeons rapidly coordinate with unfamiliar flight partners, and that familiarity has transient and relatively limited downstream effects on pair cohesion, navigational performance, and individual learning of homing routes. Examining partner familiarity is particularly tractable in small groups with repeated interactions; future work could extend this approach to investigate how learning shapes coordination in larger flocks to examine information transfer and leadership dynamics in groups with more individuals contributing to collective decision-making. Finally, our findings indicate that followership may be more repeatable and predictable than leadership in collective movement, reframing previous work in this area, and suggesting that a shift in emphasis from leadership to followership might be fruitful in improving our understanding of collective decision-making dynamics in future studies.

## Acknowledgements

This research was funded by a grant from the Templeton World Charity Foundation Inc. (TWCF-2021-20647). We would like to thank the Oxford University Biology Department for hosting the fieldwork, and Dave Wilson and Allex Turner for assisting with animal husbandry. Finally, we would like to thank Charlie Pilgrim, Elizabeth Warren, Mélisande Aellen, and all the members of the Collective Cognition Lab at the University of Rochester for their input and feedback at various stages of this project.

## Ethics statement

This work was approved by the Animal Welfare and Ethical Review Board of the Department of Biology of the University of Oxford, in accordance with University policy on the use of protected animals for scientific research, and conformed to the relevant regulatory standards.

## Data, code and materials

The dataset and code used here are available in an online repository: https://doi.org/10.5281/zenodo.21222479

## Author contributions

J.M.: Conceptualisation, Data curation, Formal analysis, Investigation, Visualisation, Writing – original draft, Writing – review & editing; P.J.L.: Conceptualisation, Investigation, Writing – review & editing; L.L.: Investigation, Methodology; G.K.: Investigation, Visualisation; J.W.K.: Investigation. T.S.: Methodology, Supervision, Writing – review & editing; R.P.M.: Data curation, Validation, Supervision, Writing – review & editing; C.K.: Funding acquisition, Supervision, Writing – review & editing; D.B.: Methodology, Funding acquisition, Supervision.

## References

Bailey, I., Myatt, J. P., & Wilson, A. M. (2013). Group hunting within the Carnivora: physiological, cognitive and environmental influences on strategy and cooperation. Behavioral Ecology and Sociobiology, 67(1), 1–17. 10.1007/s00265-012-1423-3

Baldan, D., & van Loon, E. E. (2022). Songbird parents coordinate offspring provisioning at fine spatio-temporal scales. Journal of Animal Ecology, 91(6), 1316–1326. 10.1111/1365-2656.13702

Ballerini, M., Cabibbo, N., Candelier, R., Cavagna, A., Cisbani, E., Giardina, I., Lecomte, V., Orlandi, A., Parisi, G., Procaccini, A., Viale, M., & Zdravkovic, V. (2008). Interaction ruling animal collective behavior depends on topological rather than metric distance: Evidence from a field study. Proceedings of the National Academy of Sciences, 105(4), 1232–1237. 10.1073/pnas.0711437105

Berdahl, A. M., Kao, A. B., Flack, A., Westley, P. A. H., Codling, E. A., Couzin, I. D., Dell, A. I., & Biro, D. (2018). Collective animal navigation and migratory culture: from theoretical models to empirical evidence. Philosophical Transactions of the Royal Society B: Biological Sciences, 373(1746), 20170009. doi:10.1098/rstb.2017.0009

Biro, D., Meade, J., & Guilford, T. (2004). Familiar route loyalty implies visual pilotage in the homing pigeon. Proceedings of the National Academy of Sciences, 101(50), 17440–17443.

Biro, D., Sasaki, T., & Portugal, S. J. (2016). Bringing a Time-Depth Perspective to Collective Animal Behaviour. Trends in Ecology & Evolution, 31(7), 550–562. 10.1016/j.tree.2016.03.018

Biro, D., Sumpter, D. J., Meade, J., & Guilford, T. (2006). From compromise to leadership in pigeon homing. Current Biology, 16(21), 2123–2128. 10.1016/j.cub.2006.08.087

Burt de Perera, T., & Guilford, T. (1999). The social transmission of spatial information in homing pigeons. Animal Behaviour, 57, 715–719. <GO to ISI>://WOS:000079211300024

Collet, J., Morford, J., Lewin, P., Bonnet-Lebrun, A.-S., Sasaki, T., & Biro, D. (2023). Mechanisms of collective learning: how can animal groups improve collective performance when repeating a task? Philosophical Transactions of the Royal Society B: Biological Sciences, 378(1874), 20220060. doi:10.1098/rstb.2022.0060

Couzin, I. D., Krause, J., Franks, N. R., & Levin, S. A. (2005). Effective leadership and decision-making in animal groups on the move. Nature, 433(7025), 513–516. 10.1038/nature03236

Couzin, I. D., Krause, J., James, R., Ruxton, G. D., & Franks, N. R. (2002). Collective Memory and Spatial Sorting in Animal Groups. Journal of Theoretical Biology, 218(1), 1–11. 10.1006/jtbi.2002.3065

Flack, A., Akos, Z., Nagy, M., Vicsek, T., & Biro, D. (2013). Robustness of flight leadership relations in pigeons. Animal Behaviour, 86(4), 723–732. 10.1016/j.anbehav.2013.07.005

Flack, A., Freeman, R., Guilford, T., & Biro, D. (2013). Pairs of pigeons act as behavioural units during route learning and co-navigational leadership conflicts. Journal of Experimental Biology, 216(Pt 8), 1434–1438. 10.1242/jeb.082800

Gillies, N., Tyson, C., Wynn, J., Syposz, M., Vansteenberghe, C., & Guilford, T. (2022). Exploring the mechanisms of coordinated chick provisioning in the Manx shearwater Puffinus puffinus. Journal of Avian Biology, 2022(1). 10.1111/jav.02881

Guilford, T., & Biro, D. (2014). Route following and the pigeon’s familiar area map. Journal of Experimental Biology, 217(2), 169–179. 10.1242/jeb.092908

Halliwell, C., Beckerman, A. P., Germain, M., Patrick, S. C., Leedale, A. E., & Hatchwell, B. J. (2022). Coordination of care by breeders and helpers in the cooperatively breeding long-tailed tit. Behavioral Ecology, 33(4), 844–858. 10.1093/beheco/arac048

Harcourt, J. L., Ang, T. Z., Sweetman, G., Johnstone, R. A., & Manica, A. (2009). Social Feedback and the Emergence of Leaders and Followers. Current Biology, 19(3), 248–252. 10.1016/j.cub.2008.12.051

Harcourt, J. L., Sweetman, G., Manica, A., & Johnstone, R. A. (2010). Pairs of Fish Resolve Conflicts over Coordinated Movement by Taking Turns. Current Biology, 20(2), 156–160. 10.1016/j.cub.2009.11.045

Herbert-Read, J. E., Perna, A., Mann, R. P., Schaerf, T. M., Sumpter, D. J. T., & Ward, A. J. W. (2011). Inferring the rules of interaction of shoaling fish. Proceedings of the National Academy of Sciences, 108(46), 18726–18731. 10.1073/pnas.1109355108

Kano, F., Sasaki, T., & Biro, D. (2021). Collective attention in navigating homing pigeons: group size effect and individual differences. Animal Behaviour, 180, 63–80.

Kao, A. B., Miller, N., Torney, C., Hartnett, A., & Couzin, I. D. (2014). Collective learning and optimal consensus decisions in social animal groups. Plos Computational Biology, 10(8), e1003762. 10.1371/journal.pcbi.1003762

Katz, Y., Tunstrøm, K., Ioannou, C. C., Huepe, C., & Couzin, I. D. (2011). Inferring the structure and dynamics of interactions in schooling fish. Proceedings of the National Academy of Sciences, 108(46), 18720–18725. 10.1073/pnas.1107583108

King, A. J. (2010). Follow me! I’ma leader if you do; I’ma failed initiator if you don’t. Behavioural Processes, 84(3), 671–674.

Lamprecht, J. (1991). Factors Influencing Leadership: A Study of Goose Families (Anser indicus) 1. Ethology, 89(4), 265–274.

Meade, J., Biro, D., & Guilford, T. (2005). Homing pigeons develop local route stereotypy. Proceedings of the Royal Society B: Biological Sciences, 272(1558), 17–23.

Morford, J., Gagliardo, A., Pollonara, E., & Guilford, T. (2024). Homing pigeon navigational ontogeny: no evidence that exposure to a novel release site is sufficient for learning. Anim Behav, 214, 157–164. 10.1016/j.anbehav.2024.06.009

Morford, J., Lewin, P., Biro, D., Guilford, T., Padget, O., & Collet, J. (2022). Neural networks reveal emergent properties of collective learning in democratic but not despotic groups. Animal Behaviour, 194, 151–159. 10.1016/j.anbehav.2022.09.020

Nagy, M., Ákos, Z., Biro, D., & Vicsek, T. (2010). Hierarchical group dynamics in pigeon flocks. Nature, 464(7290), 890–893. 10.1038/nature08891

Nagy, M., Vásárhelyi, G., Pettit, B., Roberts-Mariani, I., Vicsek, T., & Biro, D. (2013). Context-dependent hierarchies in pigeons. Proceedings of the National Academy of Sciences, 110(32), 13049–13054. 10.1073/pnas.1305552110

Nakayama, S., Stumpe, M. C., Manica, A., & Johnstone, R. A. (2013). Experience overrides personality differences in the tendency to follow but not in the tendency to lead. Proceedings of the Royal Society B: Biological Sciences, 280(1769), 20131724. 10.1098/rspb.2013.1724

Petit, O., & Bon, R. (2010). Decision-making processes: The case of collective movements. Behavioural Processes, 84(3), 635–647. 10.1016/j.beproc.2010.04.009

Pettit, B., Akos, Z., Vicsek, T., & Biro, D. (2015). Speed Determines Leadership and Leadership Determines Learning during Pigeon Flocking. Current Biology, 25(23), 3132–3137. 10.1016/j.cub.2015.10.044

Pettit, B., Perna, A., Biro, D., & Sumpter, D. J. (2013). Interaction rules underlying group decisions in homing pigeons. Journal of the Royal Society Interface, 10(89), 20130529. 10.1098/rsif.2013.0529

Rands, S. A., Cowlishaw, G., Pettifor, R. A., Rowcliffe, J. M., & Johnstone, R. A. (2003). Spontaneous emergence of leaders and followers in foraging pairs. Nature, 423(6938), 432–434. 10.1038/nature01630

Sasaki, T., & Biro, D. (2017). Cumulative culture can emerge from collective intelligence in animal groups. Nature Communications, 8, 15049. 10.1038/ncomms15049

Sasaki, T., Mann, R. P., Warren, K. N., Herbert, T., Wilson, T., & Biro, D. (2018). Personality and the collective: bold homing pigeons occupy higher leadership ranks in flocks. Philosophical Transactions of the Royal Society B - Biological Sciences, 373(1746). 10.1098/rstb.2017.0038

Valentini, G., Pavlic, T. P., Walker, S. I., Pratt, S. C., Biro, D., & Sasaki, T. (2021). Naïve individuals promote collective exploration in homing pigeons. eLife, 10, e68653. 10.7554/eLife.68653

Wu, C. M., Dale, R., & Hawkins, R. D. (2024). Group Coordination Catalyzes Individual and Cultural Intelligence. Open Mind, 8, 1037–1057. 10.1162/opmi_a_00155

